# The role of anionic amino acids in hydrolysis of poly-β-(1,6)-*N*-acetylglucosamine exopolysaccharides by the biofilm dispersing glycosidase Dispersin B

**DOI:** 10.1101/2020.08.05.239020

**Authors:** Alexandra P. Breslawec, Shaochi Wang, Crystal Li, Myles B. Poulin

## Abstract

The exopolysaccharide poly-*β*-(1→6)-*N*-acetylglucosamine (PNAG) is a major structural determinant of bacterial biofilms responsible for persistent and nosocomial infections. The enzymatic dispersal of biofilms by PNAG-hydrolyzing glycosidase enzymes, such as Dispersin B (DspB), is a possible approach to treat biofilm dependent bacterial infections. The cationic charge resulting from partial de-*N*-acetylation of native PNAG is critical for PNAG-dependent biofilm formation. We recently demonstrated that DspB has increased catalytic activity with de-*N*-acetylated PNAG oligosaccharides; however, there is still little known about the molecular interaction required for DspB to bind native de-*N*-acetylated PNAG polysaccharides. Here, we analyze the role of anionic amino acids surrounding the catalytic pocket of DspB in PNAG substrate recognition and hydrolysis using a combination of site directed mutagenesis, activity measurements using synthetic PNAG oligosaccharide analogs, and *in vitro* biofilm dispersal assays. The results of these studies support a model in which bound PNAG is weakly associated with a shallow anionic groove on the DspB protein surface with recognition driven by interactions with the –1 GlcNAc residue in the catalytic pocket. An increased rate of hydrolysis for cationic PNAG was driven, in part, by interaction with D147 on the anionic surface. Moreover, we identified that a DspB mutant with improved hydrolysis of fully acetylated PNAG oligosaccharides correlates with improved *in vitro* dispersal of PNAG dependent Staphylococcus epidermidis biofilms. These results provide insight into the mechanism of substrate recognition by DspB and suggest a method to improve DspB biofilm dispersal activity by mutation of the amino acids within the anionic binding surface.

## Introduction

In nature, bacteria frequently adopt a sessile lifecycle in response to environmental cues that promote the formation of surface attached biofilms (1). Biofilms consist of bacterial cells embedded in a self-assembled matrix composed of lipids, exported protein, extracellular DNA and exopolysaccharides that are known collectively as the extracellular polymeric substance (EPS) (2). The exact composition of the EPS varies depending on the bacterial species and on environmental factors, but it serves the same function in all contexts: facilitating cell–cell adhesion and acting as a protective barrier (3–6). Bacterial cells within the biofilm are shielded from the host immune response, decontamination and are often resistant to common antibiotic treatments (5, 7–9). As a result, biofilms are particularly problematic in hospital settings where biofilm formation contributes to more than half of nosocomial infections (10). Thus, approaches to either prevent biofilm formation or disrupt existing biofilms are being actively pursued to complement traditional antibiotic treatments (11–14).

Exopolysaccharides composed of poly-*β*-(1→6)-*N*-acetylglucosamine (PNAG) are a major structural constituent of biofilm EPS produced by both Gram-positive and Gram-negative human pathogens including *Staphylococcus epidermidis* (15, 16), *Staphylococcus aureus* (17) *Escherichia coli* (18), *Klebsiella pneumoniae* (19) and *Acinetobacter baumannii* (20, 21). PNAG was first identified in clinical isolates of *S*. *epidermidis* where it is commonly referred to as polysaccharide intercellular adhesin (PIA) due to its role as a primary biofilm adhesin (15). Two major forms of PNAG have been isolated from *S*. *epidermidis*. The first is a cationic polysaccharide with approximately 15% of the *N*-acetylglucosamine (GlcNAc) de-*N*-acetylated, and the second is zwitterionic as a result of both partial de-*N*-acetylation and periodic *O*-succinylation of the GlcNAc residues (16, 22). Chemical and enzymatic degradation of PNAG as well as genetic knockouts of key PNAG biosynthetic genes both result in disruption of biofilms and reduced virulence in animal infection models, highlighting the importance of this polysaccharide for biofilm integrity (8, 16, 18, 19, 23–25). Glycosidase enzymes that specifically hydrolyze PNAG have the potential to be developed as antibiofilm therapeutics as a result of their ability to disperse biofilms (11, 12, 25–27).

There have been two PNAG specific glycosidase enzymes identified to date: Dispersin B (DspB) (28) and the glycosyl hydrolase (GH) domain of the bifunctional enzyme PgaB (29). DspB is a native *β*-hexosaminidase enzyme of *Aggregatibacter actinomycetemcomitans* that has been shown to cleave PNAG using both endo- and exoglycosidic cleavage mechanisms depending on the nature of the substrate (28, 30–33). PgaB is the bifunctional carbohydrate esterase/glycosyl hydrolase enzyme required for PNAG biosynthesis in Gram-negative bacteria (24, 34). The PgaB GH domain catalyzes endoglycosidic cleavage of partially de-*N*-acetylated PNAG substrates containing a glucosamine (GlcN) in the –3 binding site (29, 35). Despite interest in these enzymes as biofilm dispersal agents and as treatments for biofilm-dependent infections, there is still relatively little known about the specific binding interactions required for recognition of their respective PNAG substrates. This is particularly true for DspB. Efforts to engineer more catalytically active variants of DspB as antibiofilm therapeutics would benefit from detailed information about the specific interactions that contribute to substrate recognition and turnover.

In a recent study, we showed that de-*N*-acetylation of PNAG oligosaccharide analogs influences both the mechanism and rate of hydrolysis of DspB for synthetic substate analogs (ie. **1** and **2**) (33). Specifically, we found that trisaccharide **2** containing GlcN at the +2 binding site showed a nearly 3-fold faster rate of exoglycosidic cleavage when compared to fully acetylated trisaccharide analog **1**. These results indicate that the substrate cationic charge may contribute to substrate recognition by DspB through interactions with anionic amino acids. Here we use a combination of site-directed mutagenesis, enzyme activity assays with synthetic PNAG substrate analogs, and *in vitro* biofilm dispersal measurements to test this hypothesis and identify amino acids involved in PNAG substrate recognition. These results suggest that mutations outside the DspB catalytic pocket influence PNAG hydrolysis activity and can be used to improve the dispersal of PNAG dependent *S*. *epidermidis* biofilms by DspB.

## Results

### Cationic charge increases the rate of substrate cleavage by DspB

Our recent studies of DspB activity using synthetic PNAG trisaccharide analogs with defined acetylation patterns (**1** and **2**) revealed that specific de-*N*-acetylation patterns influence the rate of substrate hydrolysis by DspB (33). Specifically, hydrolysis of the cationic substrate **2** was nearly 3-fold faster than for fully acetylated substrate **1**. These results suggest a hypothesis that the increased hydrolysis of **2** may result from specific charge-charge interaction with the cationic GlcN in the +2 binding site (Figure 1A). To test this further, a substrate analog **3** containing glucose (Glc) at the +2 site was synthesized. The acetylation pattern of **3** is the same as that of analog **2** but lacks the cationic charge at the +2 binding site. Analog **3** was synthesized using a one-pot sequential glycosylation approach developed for the synthesis of **1** and **2** (33), as described in detail in the supporting information.

**Figure 1.**
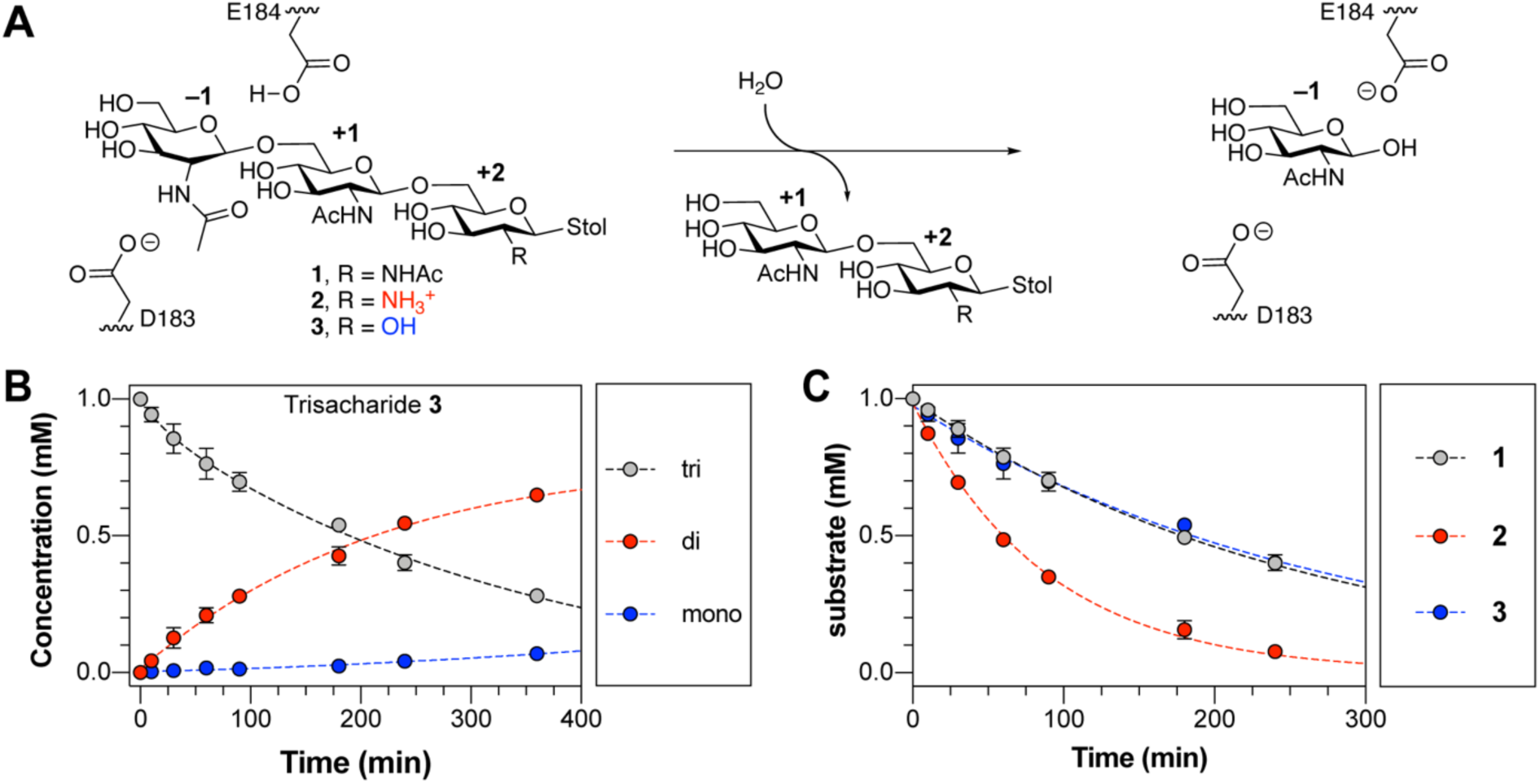
Hydrolysis of PNAG analogs by DspB. **A**. Reaction catalyzed by DspB showing the major exoglycosidic cleavage activity for trisaccharide analogs **1**–**3** used in this study. The monosaccharide residues are numbered relative to the site of glycosidic bond cleavage. The position of E184, which serves as a general acid to protonate the leaving group oxygen, and D183, which acts as a base to stabilize the oxazolinium ion intermediate, are shown. **B**. Reaction progress curve for the hydrolysis of trisaccharide **3** by DspB. Lines were added to aid identification of the disappearance of the trisaccharide (grey) and appearance of reducing-end disaccharide (red) and reducing-end monosaccharide (blue) products from sequential exoglycosidic cleavage of **3**. Error bars represent the standard deviation from at least two replicate experiments. **C**. Relative rates of trisaccharide disappearance for hydrolysis of analogs **1**–**3** by DspB. The rate of trisaccharide disappearance was fit to a single exponential using eq. 1. Error bars represent the standard deviation from two replicate experiments.

Reaction progress curves for the hydrolysis of **3** by DspB were determined by analyzing the reducing-end product distribution by HPLC (33). This assay allows for quantification of the remaining substrate and reducing-end products resulting from hydrolysis of **3** based on relative HPLC peak areas using the absorbance of the *S*-tolyl aglycone at 254 nm (Figure 1B). The reaction progress curves for hydrolysis of **3** is consistent with sequential exoglycosidic cleavage of the trisaccharide and is consistent with the mechanism observed previously for hydrolysis of **1** and **2** (33). The observed rate of hydrolysis (*k*_obs_) was determined by fitting the curve for disappearance of the trisaccharide substrate as a function of time to a single exponential using eq. 1 where [E_o_] is the initial enzyme concentration, [S_o_] is the initial trisaccharide concentration and [S] is the trisaccharide concentration remaining at time *t*.

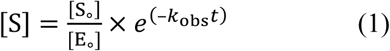

The rate of hydrolysis of **3** was nearly identical to that of **1**, and 3-fold lower than the rate of hydrolysis of cationic analog **2** (Figure 1C). This supports the hypothesis that the increased hydrolysis rate observed with **2** results from recognition of the cationic charge of the substrate, and not simply as a result of deacetylation. To further test this hypothesis and determine the specific interactions responsible for recognition of the cationic PNAG analogs, we analyzed the structure of DspB in greater detail.

### Negatively charged groove of DspB predicted to bind PNAG

A crystal structure for the DspB apoprotein was reported in 2005 (36), but structural information regarding substrate binding is lacking. DspB is classified as a Family 20 glycosyl hydrolase according to the Carbohydrate Active enZYmes (CAZy) database (37), and adopts a (β/α)_8_ TIM barrel fold, where the predicted catalytic site is found within a pocket ∼13 Å deep in the center of the *β*-barrel (Figure 2A) (30, 36). Previous GH20 enzymes have been shown to use a substrate assisted cleavage mechanism in which the oxygen of the substrate *N*-acetamido group acts as the nucleophile resulting in the formation of an oxazolinium ion intermediate (38). The GlcNAc residue at the site of bond cleavage, the –1 site, adopts an ^4^E conformation and is contained in a “cage” of conserved aromatic amino acids that serves to orient the *N*-acetamido oxygen for nucleophilic attack (38, 39). Two acidic amino acids flank the glycosidic bond, serving as a general acid protonating the leaving group oxygen (E184 in DspB) and as a “base” to stabilize the oxazolinium ion intermediate (D183 in DspB) (30, 40). The amino acids in the catalytic site are highly conserved amongst GH20 orthologs, but DspB shares little sequence conservation in the shallow binding surface surrounding the catalytic pocket (Figure 2B) (41). This is not surprising, as the GH20 family contains *β*-hexosaminidase enzymes that have activity on a range of substrates, from chito-oligosaccharides (39) to gangliosides (42) and mammalian *N*-glycan (43, 44). The only GH20 enzymes with confirmed specificity for the *β*-(1→6)-linked GlcNAc of PNAG are the DspB proteins from *A*. *actinomycetemcomitans* and *Actinobacillus pleuropneumoniae* (28, 45).

**Figure 2.**
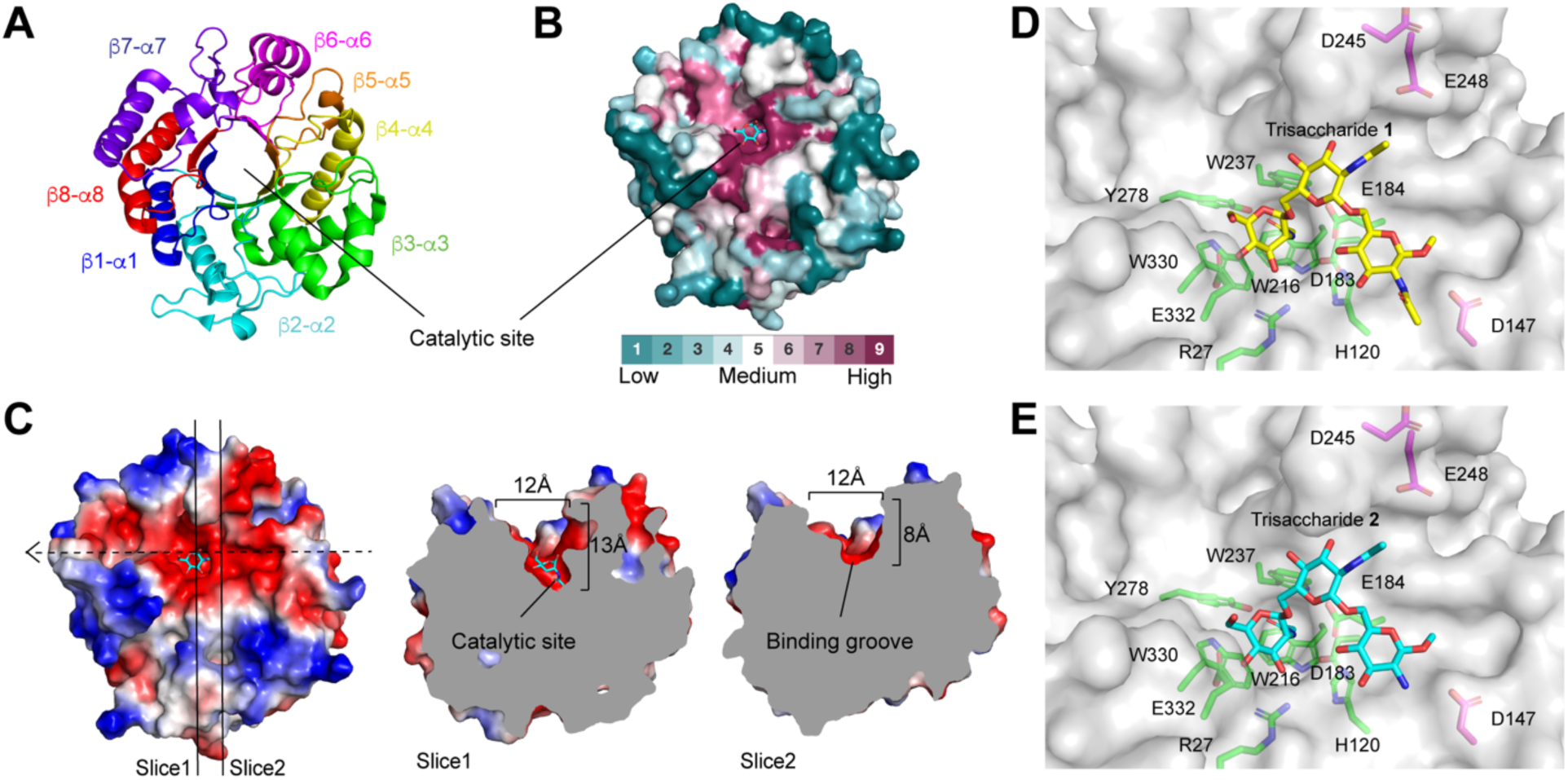
Analysis of the DspB substrate binding surface. **A**. Structure of DspB (1HYT) showing the (β/α)_8_ TIM barrel fold with each β-sheet and α-helix pair highlighted. The position of the putative catalytic site is indicated. **B**. Surface representation of DspB structure where the residues are colored based on sequence conservation amongst GH20 orthologs as calculated by Consurf (41). Cyan, white and fuchsia indicate regions of low, medium and high sequence conservation, respectively. The position of GlcNAc in the –1 site is shown in cyan sticks. **C**. Electrostatic surface map of DspB highlighting a shallow 12 Å wide anionic groove on the protein surface (indicated by dashed arrow). Two slices are shown that highlight the approximate dimensions of the binding groove. **D–E**. Low energy conformations of the methyl aglycones of trisaccharide **1** (D) and trisaccharide **2** (E) bound to DspB.

Analyzing the electrostatic surface charge of DspB identifies a number of negatively charged amino acids contributing to a shallow anionic groove adjacent to the catalytic pocket (Figure 2C). Three residues in particular, D147, D245, and E248, are located along this anionic groove and are within ∼15 Å of the catalytic site, suggesting a possible role in the recognition of cationic PNAG analogs. D147 is located in the loop connecting the β3-sheet and *α*3-helix and is conserved as an anionic amino acid (Asp or Glu) in the GH20 orthologues that are most similar in sequence to DspB. The remaining two residues, D245 and E248, are located within an extension of the *α*6-helix that is unique to DspB structure and absent in all other GH20 enzymes crystallized to date (supporting information Figure S1).

### In silico docking simulations support electrostatic protein-substrate interactions

Further support for the predicted anionic substrate binding surface was obtained from rigid body docking simulations of DspB binding to methyl glycosides of trisaccharides **1** and **2** performed using Autodock Vina (46). The highest scoring docked structure for both **1** and **2** (Figure 2D, E) adopt a nearly identical conformation in which the non-reducing end GlcNAc residue is productively positioned in the catalytic pocket in close proximity to D183 and E184. This predicted binding mode places the 2-NAc or 2-NH_3_^+^ group of the residue at the +2 site of **1** and **2**, respectively, within ∼3.2 Å of the carboxylate oxygen of D147. This supports the hypothesis that this amino acid recognizes cationic substrates through an electrostatic interaction.

### Mutation of anionic amino acids reveals their functional role in substrate recognition

To evaluate the role of anionic amino acids on PNAG substrate recognition and turnover, we mutated residues D147, D245 and E248 to the corresponding asparagine or glutamine residue or to an alanine. We also analyzed the activity of a D183A catalytic site mutant that has been previously shown to be inactive for hydrolysis of colorimetric PNAG substrate analogs (30, 47). The effect of these mutations on DspB specificity was evaluated by analyzing reaction progress curves for the breakdown of synthetic PNAG trisaccharide analogs **1**–**3** as a function of time. As with our previous studies (33), the reactions were monitored by HPLC using the absorbance at 254 nm of the *S*-tolyl aglycone to quantify the concentration of remaining substrate and all reducing end products. Obtaining steady state kinetics parameters for the DspB mutants was not possible, as the *K*_M_ values for all substrates were all >5 mM and could not be directly determined due to the limited solubility of **1**–**3** (*data not shown*). Instead, we directly analyzed reaction progress curves for the enzymatic reaction measured at a single substrate concentration of 1 mM that is well below *K*_M_ for all the mutants. The observed rate *k*_obs_ for trisaccharide hydrolysis was determined by fitting the concentration of remaining trisaccharide substrate as a function of time to a single exponential decay using eq. 1, as summarized in Figure 3A–C. Under conditions where the [S]>>*K*_M_, the reaction velocity (*v*) can be accurately described by eq. 2, where the enzyme specificity constant *k*_cat_/ *K*_M_ is equal to the observed rate constant *k*_obs_, assuming that no substrate or product inhibition is observed (48).

**Figure 3.**
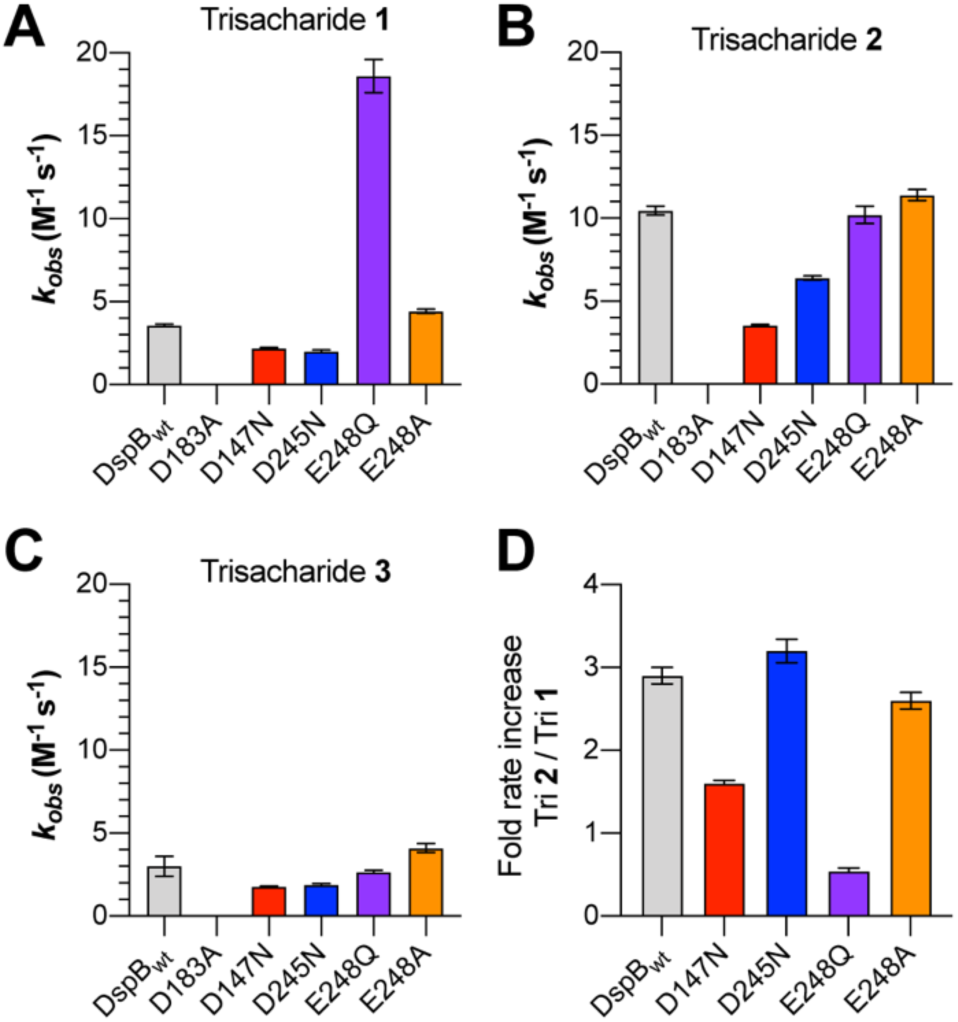
Observed rate of PNAG analog hydrolysis by DspB mutants. **A–C**. Observed rate constants (*k*_obs_) for the hydrolysis of trisaccharide **1** (A), **2** (B) and **3** (C) by DspB mutants determined by fitting the reaction progress curves for the disappearance of trisaccharide substrate to eq. 1. Error bars represent the standard deviation from two replicate time course measurements. **D**. Plot of the relative rate with cationic trisaccharide **2** compared to fully acetylated trisaccharide **1**.

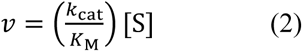

As all of the amino acid substitutions tested in this study, with the exception of D183A, occur outside of the catalytic pocket of DspB, we can assume they will predominantly influence the substrate *K*_M_. Thus, for the purpose of this study, it is assumed that the pseudo first-order rate constant *k*_obs_ is proportional to substrate binding affinity. As seen in Figure 3A–C, both the D147N and D245N mutants resulted in a slight decrease in catalytic activity (≤ 30%) when neutral trisaccharide analogs **1** and **3** were used as substrates. A nearly 3-fold decrease in activity was observed for the D147N mutant with cationic analog **2**, whereas the D245N mutant showed only a small decrease (≤ 30%) relative to DspB_wt_. This difference can be seen most clearly when we look at the relative reaction rate of the mutants with trisaccharide **2** over the rate with **1** (Figure 3D). Both DspB_wt_ (2.9 ± 0.1) and the D245N mutant (3.2 ± 0.1) display a ∼3-fold increase in rate with **2** compared to **1**, whereas the increase is only 1.6 ± 0.1 fold for the D147N mutant. This observation is consistent with a role of D147, but not D245, in recognition of cationic PNAG substrates.

The activity of DspB E248Q with cationic analog **2** or Glc containing analog **3** were statistically indistinguishable from those of DspB_wt_, but showed a 5-fold increase in catalytic activity with fully acetylated trisaccharide analog **1**. This was rather unexpected, as our hypothesis was that the mutation of anionic amino acids would predominantly affect DspB activity with cationic substrates. Since the enhanced activity was only observed for D248Q with fully acetylated trisaccharide **1**, it may result from additional hydrogen bonding between the substrate *N*-acetamido group and the amide side chain of E248Q. To test this hypothesis further, an E248A mutant was prepared. Both the activity and specificity of DspB E248A were indistinguishable from those of DspB_wt_ with all substrates tested (Figure 3A–C). This data supports our hypothesis that interactions with the amide side chain of E248Q contributes to the enhanced activity observed with analog **1**.

### Role of anionic residues in biofilm dispersal

The substrate specificity measurements reported in Figure 3 were obtained using synthetic PNAG substrate analogs and may not accurately represent the role of these anionic amino acids in the recognition of native PNAG polysaccharides. To test this, we evaluated the D147N, D245N, D248A, and D248Q mutants for their ability to hydrolyze PNAG in an *in vitro* model of *S*. *epidermidis* biofilm dispersal. *S*. *epidermidis* RP62A is a methicillin resistant isolate that produces robust PNAG-dependent biofilms on abiotic surfaces (16, 22, 49, 50). Here, biofilms of *S*. *epidermidis* RP62A that were grown in static culture for 24 hours in a 96-well microtiter plate were treated with increasing concentrations of each DspB mutant and the remaining adherent biofilm biomass was quantified by crystal violet staining (Figure 4A)(51, 52). After treating the biofilms for 90 min with DspB_wt_, there was a quantifiable reduction in adherent biofilm biomass that was dependent on enzyme concentration. Under these conditions, a biofilm dispersal EC_50_ of 240 ± 50 pM was measured for DspB_wt_ (Figure 4B). Treating with the catalytically inactive D183A mutant resulted in less than 20% dispersal, even after 90 min treatment with 2.5 *μ*M enzyme, which is consistent with this residue’s role in stabilizing formation of the oxazolinium ion intermediate during PNAG hydrolysis (30). As seen in Figure 4B, the EC_50_ for the D147N mutant (480 ± 100 pM) was nearly 2-fold greater than that of DspB_wt_ while those for D245N (320 ± 85 pM) and E248A (280 ± 110 pM)

**Figure 4.**
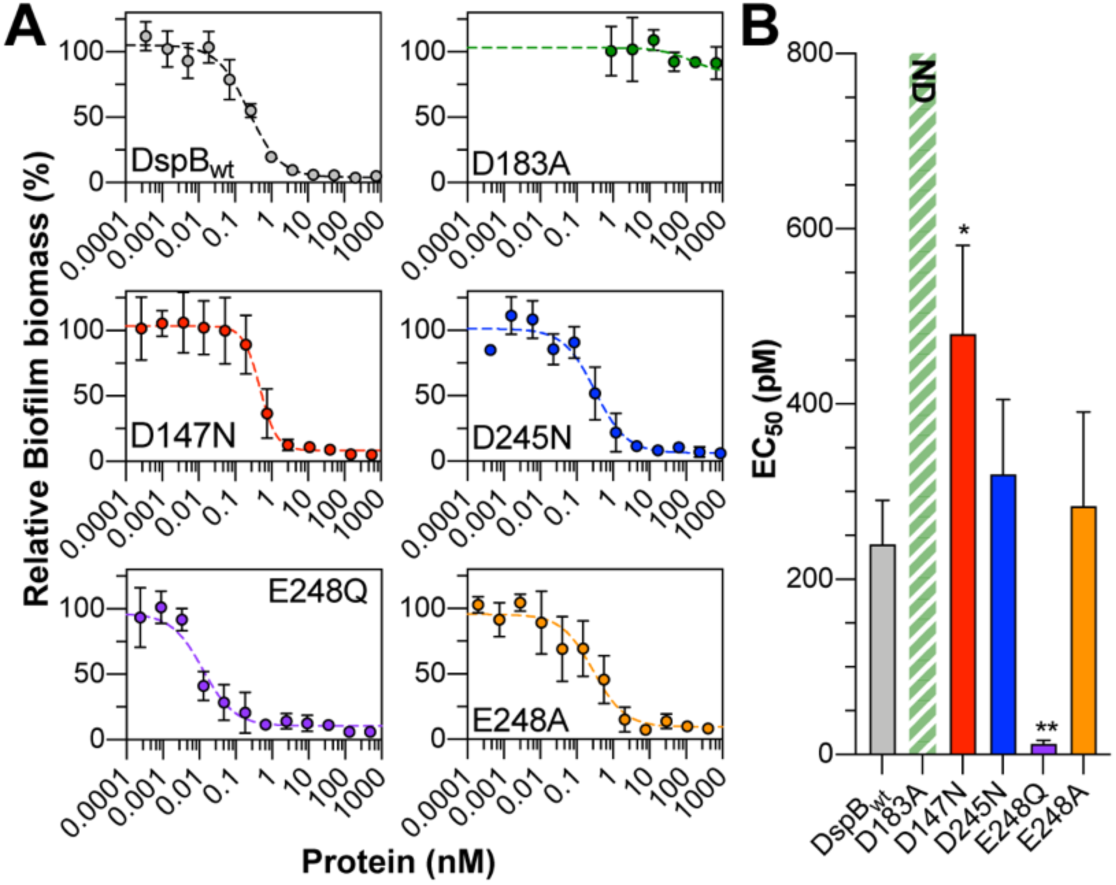
Dispersal of S. epidermidis biofilms by DspB mutants. **A**. Plots of *S*. *epidermidis* RP62A biofilm biomass remaining after treating with varying concentrations of DspB mutants for 90 min relative to untreated biofilms. Error bars represent the standard deviation of four biological replicates. **B**. EC_50_ values from *S*. *epidermidis* biofilm dispersal assays. Error bars represent the standard deviation from four biological replicates. Statistical significance as compared to DspB_wt_ was determined using a one-way ANOVA with Dunnett’s multiple comparison. *, P < 0.05; **, P < 0.01. ND: the EC_50_ was not determined as > 75 % biofilm biomass remained even after treating with 2.5 μM enzyme for 90 min.

were the same as DspB_wt_, within experimental error. This is consistent with the observed activities for these mutants with trisaccharide analogs **1**–**3**. The E248Q mutant had the lowest EC_50_ at 13 ± 4 pM, consistent with the enhanced catalytic activity observed for this mutant with fully acetylated PNAG analog **1**.

### Discussion

Given the importance of PNAG for biofilm integrity and its widespread distribution amongst Gram-positive and Gram-negative bacteria, enzymes that catalyze the hydrolysis of PNAG are potentially useful for treating diverse biofilm infections (11, 12). Two PNAG glycosidases, DspB and the *C*-terminal domain of PgaB, have been described to date, but both suffer from relatively low catalytic activity (29, 30, 35). For example, the activity of DspB measured with a variety of substrate analogs has resulted in observed rates in the order of 2–60 M^-1^s^-1^ (30, 32) nearly 4–6 orders of magnitude slower than comparable rates observed with other GH20 enzymes (40, 53). The poor catalytic activity of DspB likely results from the low binding affinity between DspB and its PNAG substrate. In fact, measurements of DspB binding affinity for PNAG oligosaccharides of varying lengths resulted in measured dissociation constants (*K*_d_) of between 1–10 mM (54). Efforts to improve DspB catalytic efficiency would benefit from a detailed analysis of substrate binding.

Our previous studies showed that DspB has nearly 3-fold greater catalytic activity with substrate analog **2** containing GlcN in the predicted +2 binding site compared to a fully acetylated analog **1** (33). In the studies described here, we confirmed this substrate preference and demonstrated that it is the cationic charge of GlcN at the +2 site that is responsible for the increased catalytic activity. This suggests that anionic amino acid residues of DspB contribute to PNAG substrate binding. The results support PNAG binding along a shallow anionic groove on the surface of DspB with D147 contributing to the recognition of cationic GlcN residues in the PNAG substrate. A relatively modest decrease in activity was observed for a D147N mutant, consistent with its role in recognition of cationic PNAG substrates, but also indicating that other residues along the substrate binding surface likely contribute to the recognition of cationic PNAG substrates as well. The reduced activity of DspB D147N with synthetic substrate analogs was consistent with observed biofilm dispersal activity of PNAG-dependent *S*. *epidermidis* biofilms in a surface attached biofilm model.

Neither of the other anionic amino acids, D245 and E248, appear to contribute to the specificity of DspB for trisaccharide analogs **1**–**3** or to the *in vitro* dispersal of *S*. *epidermidis* biofilms, as D245N and E248A mutants had activity that was indistinguishable from DspB_wt_ in these assays. These results are consistent with the bound conformation of trisaccharide **1** and **2** predicted from rigid body docking simulations (Figure 2D and E). In these models, the non-reducing GlcNAc residue occupies the catalytic pocket in an orientation consistent with the crystal structures of related GH20 enzymes, including *Ostrinia furnacalis* Hex1 (55), *Serratia marcescens* chitobiase (39) *Bifidobacterium bifidum* lactobiase (56), *Streptococcus pneumoniae* StrH (44), and *Streptomyces plicatus* β-*N*-hexosaminidase (57). Compared to these GH20 enzymes, the surface surrounding the catalytic pocket of DspB is shallow and lacks aromatic amino acid residues that contribute to substrate recognition through C-H/*π* interactions that are present in most of the other GH20 enzymes (Figure S1). The docked conformations of both **1** and **2** place the 2-NHAc or -NH_3_ ^+^ group of the +2 residue within 3.2 Å of the carboxylate oxygen of D147. Such a conformation would allow for either hydrogen bonding, in the case of **1**, or the formation of a charge-charge interaction, in the case of **2**, consistent with the reduced catalytic activity observed for the D147N mutant in this study.

Significant interactions were observed between amino acids in the catalytic pocket and the substrate –1 GlcNAc residue and between D147 and the +2 residue, but there was a lack of other stabilizing interactions observed between DspB and the +1 and+2 residues of both docked trisaccharides **1** and **2**. The lack of specific binding interactions helps to explain the relatively poor binding affinity that has been observed between DspB and PNAG analogs *in vitro* (54). Interestingly, the E248Q mutant, located within the *α*6-helix extension that is unique to the structure of DspB, had no impact on activity with cationic substrate analog **2**, but resulted in a 5-fold increase in activity with fully acetylated analog **1**. The same activity was not observed for the neutral deacetylated analog **3**, supporting the hypothesis that the E248Q mutation introduces new interactions unique to the fully acetylated PNAG substrate. The increased activity was not observed with E248A that lacks the side chain amide, supporting a role for hydrogen bonding between the side chain amide of E248Q and the 2-acetamido group at the +2 site of trisaccharide **1**. Such an interaction was not observed in the docked structure of **1** (Figure 2D) that orients the +2 GlcNAc more than 9 Å from E248. It is important to note that these in silico docking simulations were performed using the DspB apo structure (PDB 1YHT) (36). It is possible that substrate binding induces a conformational change in DspB that would place E248Q in closer proximity to the bound substrate. This conformational flexibility of the *α*6-helix extension is supported, at least in part, from an analysis of B-factors for the DspB apo structure (36). The *α*6-helix extension and residues in the anionic binding surface have among the highest B-factors (Figure S2) indicating these residues reside in an area of increased flexibility in the DspB apo structure. Further structural studies of DspB with PNAG substrate analogs are required to analyze this flexibility in greater detail.

Taken together, the results of the studies presented here support a model in which PNAG binds along a shallow anionic groove on the DspB surface, and D147 contributes to the recognition of cationic PNAG substrates through charge-charge interactions. Neutral, fully *N*-acetylated PNAG likely adopts multiple unique bound conformations within the substrate binding site, and this binding is driven by interactions with GlcNAc at the –1 binding site. Flexibility in the PNAG binding site would accommodate substrates with diverse and heterogeneous PNAG modifications, such as percent de-*N*-acetylation and succinylation, that are known to vary depending on the bacterial species and environmental factors (16, 22). Moreover, we showed that site-directed mutagenesis of residues lining this anionic binding surface can alter the catalytic turnover of PNAG substrate analogs such as **1**–**3**, and that mutations that improve the catalytic efficiency with oligosaccharide substrates correlate with improved dispersal of PNAG dependent biofilms *in vitro*. Future work should focus on mutations of the residues lining the anionic binding surface to engineer DspB mutants that may function as more effective biofilm dispersal agents for treatment of diverse biofilm-dependent infections.

## Experimental Procedures

### Protein production

Recombinant Dispersin B (residues 16-381, DspB_wt_) from *A*. *actinomycetemcomitans* was prepared as described previously (33). Plasmids for expression of DspB_D183A_ were prepared as previously described (47). Plasmids for expression of DspB_D147N_, DspB_D245N_, DspB_E248Q_, and DspB_E248A_ mutants were prepared by Quikchange site-directed mutagenesis using the primer pairs outlined in Table 1. All mutations were confirmed by single-pass Sanger sequencing and were expressed recombinantly in *E*. *coli* BL21(DE3) and purified as described for DspB_wt_ (33). Proteins were quantified by UV absorbance at 280 nm using a calculated molar extinction coefficient of 51,340 M^−1^cm^−1^. All proteins were purified to > 95% purity as confirmed via SDS-PAGE (Figure S3), flash frozen, and stored as individual aliquots at –80°C for later use.

**Table 1.**
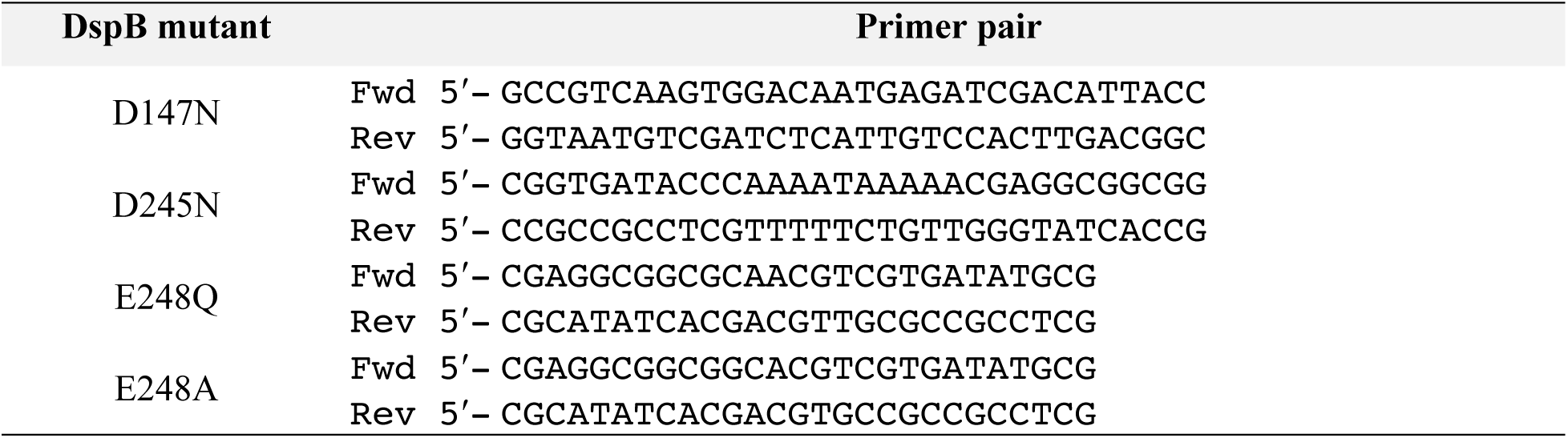
Primers for site directed mutagenesis used in this study.

### Rigid body docking simulations

Trisaccharide substrate models for the methyl glycosides of **1** and **2** were prepared with the non-reducing GlcNAc residue adopting a ^4^E ring conformation observed in crystal structures of related GH20 enzymes with bound substrate (39, 44, 55, 57). Models for the rest of the trisaccharide substrates were prepared using the lowest energy ^4^C_1_ ring conformers and glycosidic bond conformations generated using glycan builder on the glycam.org webserver (58), and prepared using in Open Babble. Ring conformations were fixed while all other bonds were allowed to freely rotate. Rigid body docking simulations for the binding of **1** or **2** were carried out using Autodock Vina (46). The highest-scoring ligand conformers for both trisaccharides adopted the same conformation with the non-reducing terminal GlcNAc residue bound in the catalytic pocket in the same orientation as observed in other GH20 enzymes. These conformers were exported for figure preparation in Pymol (version 2.3.0).

### Time course assays for hydrolysis of PNAG analogs 1–3

Reaction progress curves for hydrolysis of analogs **1**–**3** by DspB mutants were measured by HPLC using the previously reported approach with a few modifications (33). Briefly, hydrolysis reactions containing 1 mM of trisaccharide (**1–3**) in 48 mM potassium phosphate, pH 6.0 buffer containing 100 mM NaCl were initiated by the addition of an appropriate concentration of DspB_wt_ or DspB mutant enzyme in a final volume of 50 *μ*L. Individual 5 *μ*L aliquots were removed after 0 min, 10 min, 30 min, 60 min, 90 min, 180 min, 240 min and 360 min incubations at 22 °C and quenched through the addition of 5 *μ*L of 100 mM trifluoroacetic acid (TFA). Quenched fractions were centrifuged at 17,000 × g for 2 min to pellet any insoluble material, diluted to 50 *μ*L with MQ water, and analyzed by reversed phase HPLC as described previously (33). The concentration of residual substrate, and all reducing-end products were determined from their relative peak areas based on the absorbance at 254 nm resulting from the *S*-tolyl aglycone. Reaction rates for trisaccharide hydrolysis were determined by plotting the concentration of residual trisaccharide as a function of time and fitting to a single exponential using eq. 1 where [S] is the trisaccharide concentration at time *t*, [S_o_] is the initial trisaccharide concentration, and [E_o_] is the initial enzyme concentration. This gives a normalized pseudo-first order rate constant *k*_obs_ in units of M^−1^s^−1^.

### *S*. *epidermidis* biofilm dispersal assays

*S*. *epidermidis* RP62A was obtained from ATCC (ATCC^®^ 35984™). For biofilm dispersal assays, a starter culture was grown in 25 mL of tryptic soy broth (TSB) for 24 h at 37 °C with shaking. The starter culture was then diluted to an OD_600_ of 0.01 using sterile TSB and 200 *μ*L of the diluted culture was added to each well of a clear, flat-bottom 96-well plate (ThermoFisher Nunc™ Edge™). MQ water was added to the outer moat according to the manufactures recommendation to limit evaporation and edge effects during growth. The plates were grown in static culture for 24 h at 37 °C. After 24 h, the culture media and all non-adherent cells were removed by aspiration and 200 *μ*L of sterile 50 mM potassium phosphate buffer, pH 6.0 was added to each well. 20 *μ*L of a solution containing various dilutions of DspB_wt_ or mutant enzyme in 50 mM potassium phosphate buffer, pH 6.0 was added to the wells to a final volume of 220 *μ*L. A minimum of eight replicates on each plate were incubated with buffer alone and functioned as no dispersal controls (0% dispersal). The plates were incubated with enzyme at 25 °C for 90 min, at which time the buffer solution was removed by aspiration and the plates were washed gently but thoroughly with DI water to remove any non-adherent biomass. Remaining adherent cells were fixed with MeOH (200 *μ*L) for one hour after which the wells were aspirated and allowed to fully dry. This was followed by staining with 1% crystal violet (200 *μ*L) for 5 min. The wells were rinsed thoroughly with DI water until the water ran clear. The plates were imaged to document the stained biofilm biomass.

To quantify the adherent biomass, 200 *μ*L of 33% acetic acid was added to each well to release the crystal violet, and 50 *μ*L was removed and diluted with an additional 150 *μ*L of 33% acetic acid in a separate 96 well micro-titer plate. The absorbance of crystal violet in each well was measured at 590 nm using a plate reader. The relative biofilm dispersal was calculated by dividing the absorbance at 590 nm by the average absorbance of the no dispersal control wells from each plate. The relative biofilm biomass was plotted as a function of enzyme concentration to determine EC_50_ values for biofilm dispersal. Each enzyme concentration was analyzed in quadruplicate in different positions on the 96-well plate to minimize artifacts from edge effects. Statistical significance of EC_50_ values relative to DspB_wt_ were determined using a one-way ANOVA with Dunnet’s multiple comparison as implemented in GraphPad Prism 8 software.

## Data availability

All data supporting the findings in this study are available within the article and its supporting information.

## Author Contributions

A.P.B. and C.L. produced all proteins, and A.P.B. carried out all activity and in vitro biofilm dispersal assays. S.W. synthesized trisaccharide substrates **1**–**3**. A.P.B. and M.B.P. wrote the manuscript. All authors analyzed data and reviewed the final version of the manuscript.

## Acknowledgments

We are grateful to F. Chen and Y. Li for assistant with analytical NMR and mass spectrometry services.

## Funding and additional information

This work was funded in part by startup funds and a Faculty Student Research Award from the Graduate School at University of Maryland College Park.

